# Leveraging large language models for metabolic engineering design

**DOI:** 10.1101/2024.09.09.612023

**Authors:** Xiongwen Li, Zhu Liang, Zhetao Guo, Ziyi Liu, Ke Wu, Jiahao Luo, Yuesheng Zhang, Lizheng Liu, Manda Sun, Yuanyuan Huang, Hongting Tang, Yu Chen, Tao Yu, Jens Nielsen, Feiran Li

## Abstract

Establishing efficient cell factories involves a continuous process of trial and error due to the intricate nature of metabolism. This complexity makes predicting effective engineering targets a challenging task. Therefore, it is vital to learn from the accumulated successes of previous designs for advancing future cell factory development. In this study, we developed a method based on large language models (LLMs) to extract metabolic engineering strategies from research articles on a large scale. We created a database containing over 29006 metabolic engineering entries, 1210 products and 751 organisms. Using this extracted data, we trained a hybrid model combining deep learning and mechanistic approaches to predict engineering targets. Our model outperformed traditional metabolic engineering target prediction algorithms, excelled in predicting the effects of gene modifications, and generalized well to out-of-distribution products and multiple gene combinations. Our study provides a valuable dataset, a chatbot, and an engineering target prediction model for the metabolic engineering field and exemplifies an efficient method for leveraging existing knowledge for future predictions.

## Introduction

The establishment of efficient cell factories is fundamental for synthetic biology, aiming to optimize microbial and cellular systems to produce valuable chemicals, fuels, and pharmaceuticals^1^. However, this process is inherently complex and iterative, often leading to extensive trial and error to identify effective engineering targets. Genome-scale metabolic models (GEMs) are regarded as metabolism-related knowledge database and have been instrumental in providing a framework for understanding these interactions, which can simulate cellular metabolism based on stoichiometric relationships and have been used to predict the effects of gene knockouts, over-expressions, and environmental changes^2–7^. Despite their utility, GEMs rely on simplified assumptions and lack of information, such as regulatory mechanisms, limiting their effectiveness in metabolic engineering applications. Additionally, current GEM-based algorithms typically struggle with predicting the synergistic effects of multiple gene modifications, especially with gene overexpression^5^.

The accumulating wealth of successful metabolic engineering strategies is a particularly valuable practical data resource, which has however yet to be fully exploited. Manual data extraction from numerous articles poses challenges in maintaining consistency and accuracy, requires substantial time^8, 9^. Significant advancements in natural language processing (NLP) have been made with the introduction of large language models, such as GPT-3.5^10^, GPT-4^11^, Llama-2^12^, Llama-3^13^, and Qwen1.5^14^. These models are revolutionary because they can analyze, comprehend, and generate human language on an unprecedented scale. With prompt engineering, retrieval-augmented generation (RAG) techniques, and fine-tuning with domain-specific knowledge, LLMs have shown superior performance in information extraction, deduction, and answer specialized and expert-level in scientific related tasks such as to guide chemical synthesis^15, 16^, medical research^17, 18^ and material design^19–22^.

Recent advances have shown that integrating data-driven insights into mechanistic models such as GEMs can yield impressive results, even when trained with small amounts of in-house data or a limited number of organisms cases. This approach has been successfully applied to various areas, including metabolic flux prediction^23^, transcriptome changes predictions from gene perturbations^24^, tryptophan phenotype predictions^25^, antimicrobial resistance phenotype predictions^26^, chemical yield production^27^ and bacteria growth rate predictions^23^. Drawing inspiration from these advancements, we see great potential to extract comprehensive practical data sources from previous publications and integrate with the theoretical knowledge databases such as GEMs. By leveraging data mining of accumulated metabolic engineering strategies, we can enhance our predictive capabilities and optimize metabolic pathways more efficiently. This integration could lead to improved strain design, more effective bioproduction processes, a deeper understanding of cellular metabolism and perspectives and tools for biological research.

In this study, we develop an LLM-based pipeline called D2Cell (Deep learning-aided Design of Cell factories) to extract metabolic engineering knowledge and advance the metabolic engineering prediction. We employed the D2Cell pipeline to extract information such as products, metabolic engineering strategies, product titer, and fermentation conditions. We conducted extensive manual validations to assess the model’s performance. Additionally, leveraging the collected data, we developed a hybrid model combining deep learning and GEM to predict metabolic engineering targets for product overproduction, and tested the model performance on the out-of-distribution products (those not included in the training dataset) and on the impact of multiple gene modifications. Furthermore, we fine-tuned a cell factory-related conversational chatbot to answer metabolic engineering questions for further usage.

## Results

### D2Cell design

We developed an LLM-based pipeline, D2Cell, for designing future cell factories (Fig. 1). The pipeline includes three main components: information collection, information extraction and application. For efficient extraction of engineering strategies from literature, we decomposed the complex task into simpler sub-tasks^28, 29^, enhancing LLM output accuracy. The extraction process was divided into three phases: named entity recognition (NER), relation extraction (RE), and entity resolution (ER). The D2Cell pipeline effectively created a structured cell factory database by utilizing NER to identify key entities (e.g., organisms, strain IDs), RE to link gene modifications and extract relevant conditions, and ER to ensure database quality and consistency. This process enhances the accuracy of the knowledge base and facilitates its integration with existing biological databases.

**Fig. 1.**
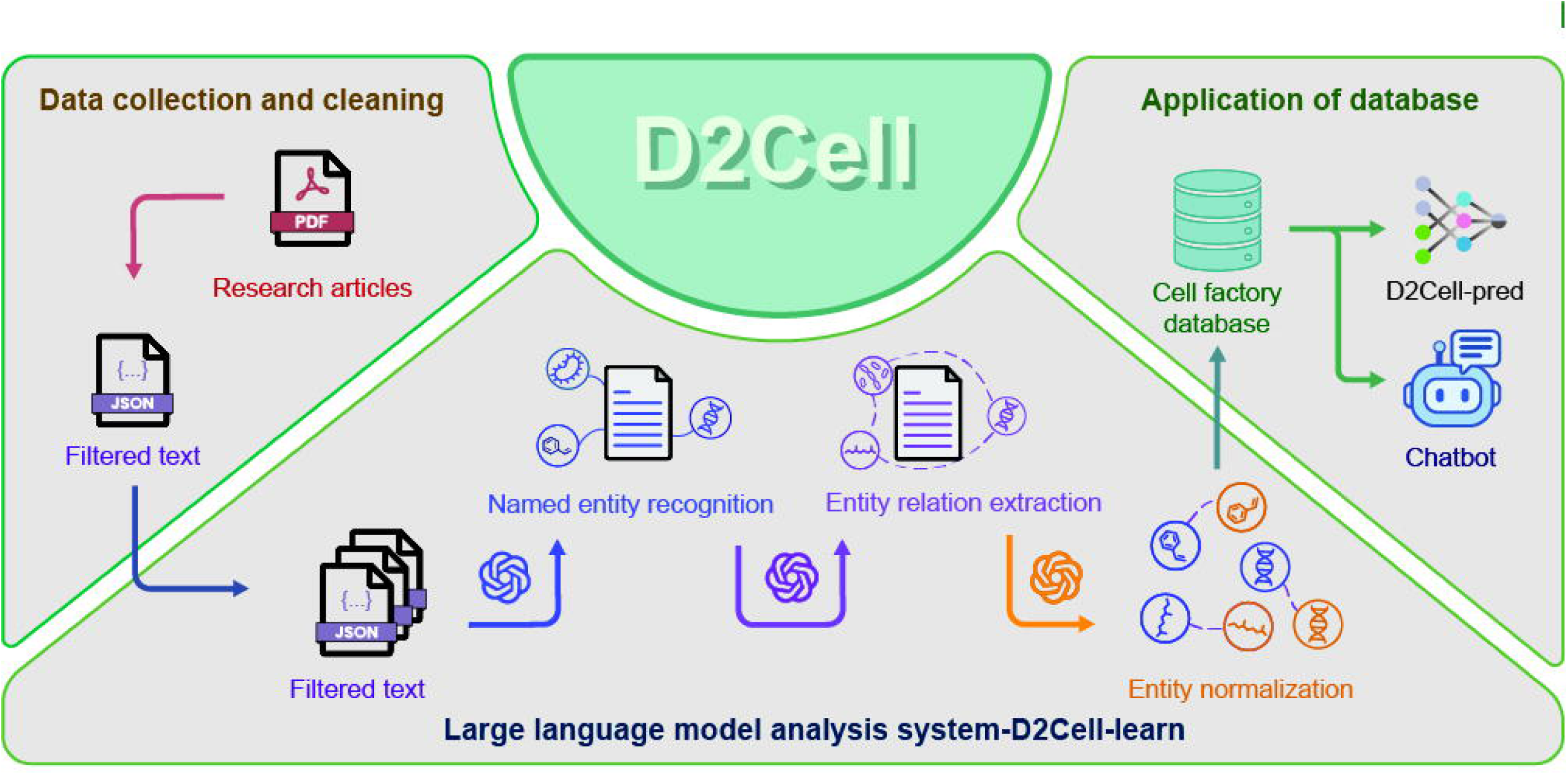
D2Cell pipeline for cell factory analysis workflow. The workflow contains three modules. Data collection and cleaning module collects and extracted filtered text from literature. The large language model analysis system (D2Cell-learn) includes named entity recognition (NER), entity relation extraction (RE) and entity normalization (ER). NER extracts key entities from preprocessed text such as organisms and strain IDs. RE extracted relationships between key terms based on the text content such as linking the titer to gene modification. ER standardizes the key entities by normalizing the naming from different literature, such as ‘D-glucose’ with ‘glc’. The extracted data were then formatted into a cell factory database. Using this database, a hybrid model D2Cell-pred combining deep neural network and mechanistic metabolic model was trained to predict genetic modification effects, and a metabolic engineering chatbot was developed.

### D2Cell development and comparison with LLMs in text mining

After designing the model structure, we evaluated various LLMs for each module, including GPT-4, Gemini Pro^30^, Claude-3^31^, Llama-3^13^, and Qwen1.5^14^. In the absence of a suitable NER dataset for microbial genes and strain IDs, we manually annotated a dataset to evaluate performance. We selected 873 text segments covering various metabolic engineering sources, which were then manually annotated and cross-validated by multiple synthetic biology researchers (Supplementary File 1). Comparison of the labeled data with the LLM-annotated results showed that GPT-4, Qwen1.5, and Llama-3 had the high recall but low precision among the base LLMs (Supplementary Fig. 1a), which indicates that these models are effective at identifying a wide range of relevant instances (high recall), but they also include a considerable number of incorrect or irrelevant information (low precision). On the other hand, Gemini Pro^30^, though not as comprehensive, showed high precision. To enhance LLMs’ performance, we manually verified data from Gemini Pro and GPT-4 to fine-tune Qwen1.5-14B-Chat and Llama3-8B-Instruct, resulting in Qwen-Lora and Llama3-Lora. The fine-tuned model exhibited significant performance enhancements in benchmarks, with substantial improvements in accuracy and F1 scores compared to the original and other models (Supplementary Table 1). The Qwen-Lora model achieved the highest precision (85%) and F1 score (86%) and maintain a high recall (86%), which indicates the Qwen-Lora model is both accurate and thorough. Qwen-Lora processed the 873 texts in just 27 minutes, making it one of the fastest (Supplementary Fig. 1b). Since maintaining the data extraction correctness is the main goal to ensure the performance in the further prediction, thus Qwen-Lora was selected for the NER task in D2Cell-learn, with GPT-4 also available for comparison.

Relation extraction (RE) identifies relationships between entities, such as linking gene modifications to products and titers. Unlike the NER task, which focuses on individual sections of a paper, the RE task requires input from the abstract, methods, and results sections, as relationships between entities often span multiple sections. As for the LLMs choice for RE task, the Llama-3 models were excluded from this step because their 8096-token limit wasn’t enough to handle full-text inputs. Instead, we selected the top-performing base models from the NER task, Qwen1.5-110B-Chat (open-source) and GPT-4 (commercial) for the RE task.

Since it’s challenging to evaluate the RE task separately from the NER component, we assessed the overall performance of D2Cell-learn, which includes both tasks. We selected a collection of 100 metabolic engineering papers for performance analysis. Two versions of D2Cell: D2Cell-Qwen (default) and D2Cell-GPT4, along with other LLMs, were utilized to extract relevant data from these publications. The extracted information included metabolic engineering strategies, organisms, products, product titers, and medium setup. Subsequently, we conducted manual verification of the extracted data to assess its accuracy. Overall, the D2Cell method achieved 74% and 79% accuracy (Fig. 2a), with a median paper-based accuracy nearing 100% (Supplementary Fig. 1c), outperforming direct extraction with Qwen1.5 and GPT-4. This highlights the benefits of dividing tasks into NER and RE. Additionally, the D2Cell models substantially increased the volume of extracted data, resulting in a 50-100% increment in data extraction compared to direct methods (Fig. 2a). This significant increase in the data extraction quantity underscores the method’s enhanced capability for comprehensive information extraction from complex textual sources. While D2Cell-GPT4 performs slightly better, it incurs higher costs (Fig. 2a). We evaluated the execution time of each module on D2Cell-Qwen (Fig. 2b). The results indicate that the RE module is the most time-intensive component within the D2Cell workflow. The significant time cost is driven both by the large number of tasks in the RE module and the complexity of each task, as they require handling large volumes of information and producing more detailed outputs. As a result, the RE module has a significantly higher time requirement compared to the other modules and its performance has a critical impact on the overall performance.

**Fig. 2.**
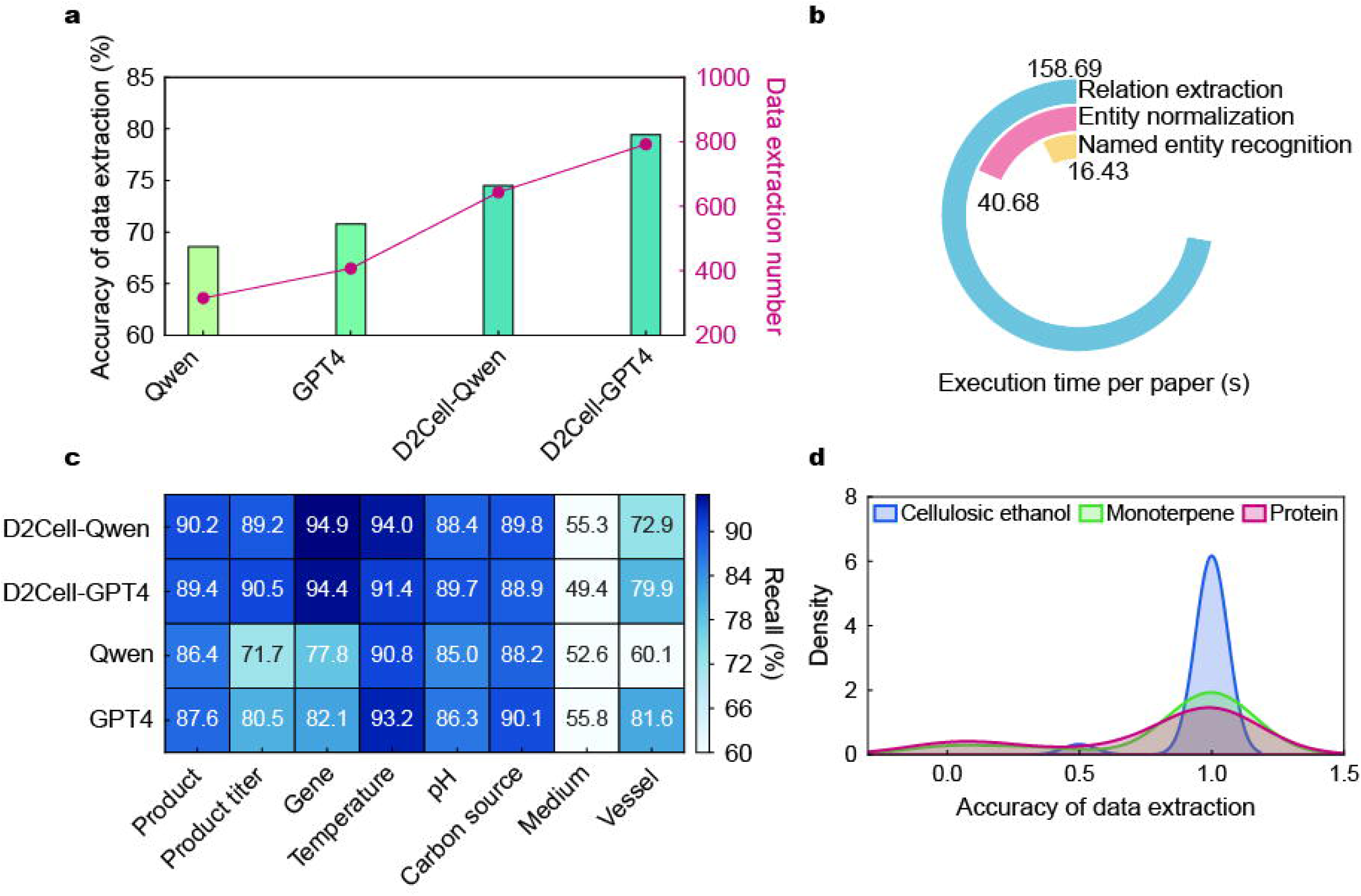
Performance analysis of D2Cell-learn. a) Accuracy and quantity of metabolic engineering data extracted from a test set of 100 cell factory articles. b) Comparison of the average execution time required by each module of D2Cell-Qwen to process a single paper. The unit is in second. c) The detailed percentage recall for each entity across the different LLMs on the LASER dataset. d) Accuracy on data extraction for three types of products of D2Cell-Qwen.

Then, we validated the model performance with the LASER dataset, an independent and manually curated test set described in a previous study^9^. Both D2Cell-Qwen1.5 and D2Cell-GPT-4 achieved the highest recall scores for most categories, outperforming directly using GPT-4 and Qwen1.5. While D2Cell faced challenges with low recall in extracting medium information due to terminology inconsistencies across papers, it excelled in accurately retrieving data on gene modifications, products, and organisms, achieving recall of approximately 90% in these three crucial entity types (Fig. 2c). Moreover, to test whether D2Cell has any preference towards product types, we assessed its performance across three main product types—cellulosic ethanol (commodity chemicals), monoterpenes (natural products), and recombinant proteins (protein-based products). Verification by synthetic biology human experts confirmed D2Cell’s high accuracy across these categories (Fig. 2d), with an average extraction accuracy approaching 1 (Supplementary File 3). Taken together, these tests demonstrate the superior performance of D2Cell and highlight that breaking down complex tasks into simpler subtasks^29^ can effectively enhance the accuracy of key information extraction from literature.

### D2Cell assisted metabolic engineering database construction

Using the D2Cell workflow, we processed the metabolic engineering literature published from 2000 to 2023 by various publisher groups, including more than 10000 abstracts and 1340 open-access full texts (Supplementary Fig. 2a). We successfully identified, extracted and standardized in total 29006 data entries. Each entry includes details on the organism, product, product titer, genetic modification, medium setup, oxygen availability, volume, and temperature. This database comprises data from 751 microbes and 1210 unique products, with *Escherichia coli* (35.0%) and organooxygen compounds (28.9%) being the most represented organism and product, respectively (Fig. 3a-b). As for the product-based analysis, we observed a significant increase in both the volume of papers and the diversity of products studied after 2009 (Fig. 3c), which indicates a substantial expansion in research and applications within metabolic engineering. Furthermore, we analyzed the product distribution across various host organisms and found that ethanol was the favorite product for *Saccharomyces cerevisiae* and *Kluyveromyces marxianus*, while lysine production was a primary focus for *Corynebacterium glutamicum* (Fig. 3d). Our analysis of metabolic engineering strategies for product synthesis in *E. coli* revealed patterns consistent with Z. A. King et al.^32^, demonstrating frequent knockdowns of phosphate acetyltransferase (pta), acetate kinase (ackA), and formate acetyltransferase (pflB) genes to enhance succinate production, while Bifunctional aldehyde-alcohol dehydrogenase (adhE) and formate acetyltransferase (pflB) gene knockdowns were commonly employed to overproduce D-lactate (Supplementary Fig. 2b). All extracted entries are available as a cell factory database (LINK). As literature accumulates and the database continues to improve, it will serve as a comprehensive resource for metabolic engineering, supporting further exploration and development in the field.

**Fig. 3.**
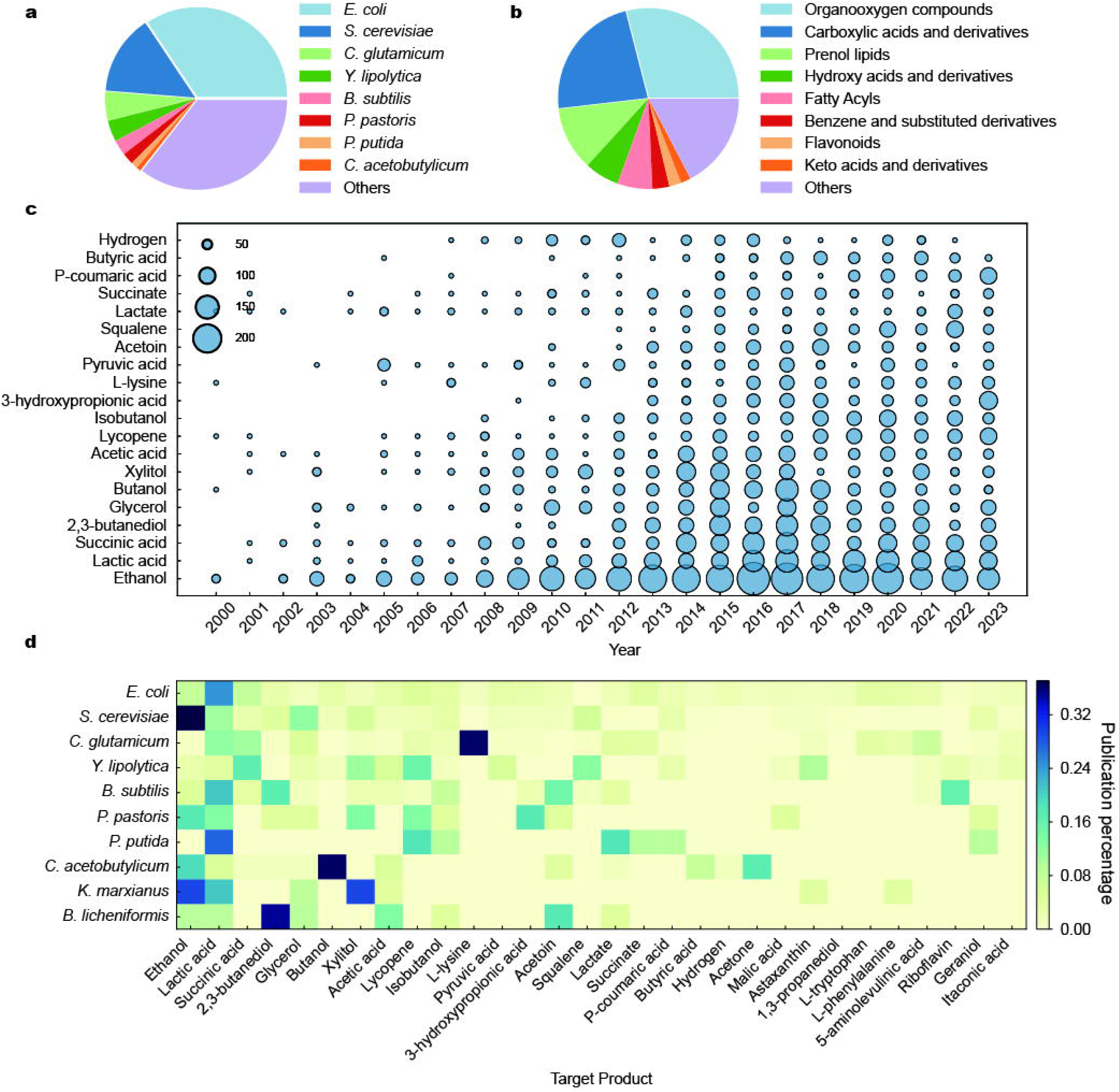
The D2Cell database. a) Distribution of organisms in the database. b) Distribution of product classes in the database. c) Number of papers published from 2000 to 2023 for the top 20 products in the database. d) Distribution of the top 30 products across the top 10 organisms. Percentage is normalized for each organism. *E. coli*: *Escherichia coli*, *S. cerevisiae*: *Saccharomyces cerevisiae*, *C. glutamicum*: *Corynebacterium glutamicum*, *Y. lipolytica*: *Yarrowia lipolytica, B. subtilis*: *Bacillus subtilis, P. pastoris*: *Pichia pastoris, P. putida*: *Pseudomonas putida, C. acetobutylicum*: *Clostridium acetobutylicum*, *K. marxianus*: *Kluyveromyces marxianus, B. licheniformis*: *Bacillus licheniformis*.

### Query-assisted conversational chatbot for metabolic engineering

To enhance the utility of our comprehensive metabolic engineering database, we developed an interactive, user-friendly conversational chatbot (LINK). Employing the Retrieval-Augmented Generation (RAG) technique^33^, the chatbot improves data accessibility by generating responses grounded in accurate information from the database. This ensures that the provided information is both relevant and derived from our literature mining process.

We first created tabular entries corresponding to products and strains in the dataset (Fig. 4). These entries include a wide range of experimental conditions—such as fermentation time, gene modifications, and carbon sources—as well as bibliographic details like authors, DOI, and publication date. We then generated embeddings from these tabular data to construct a vector dataset. The chatbot retrieves the most relevant entries from this vector dataset based on user queries. It concatenates these entries with the query to form prompts for the LLM, which then generates responses based on the retrieved data, minimizing potential hallucination errors^34^. Additionally, the chatbot provides DOI links to the original literature, facilitating direct access to sources and ensuring data traceability (Fig. 4). Furthermore, leveraging the extensive local metabolic engineering database, the chatbot offers detailed explanations and examples of complex theories, rather than vague responses (Fig. 4). This design enhances the chatbot’s ability to elucidate concepts and experimental procedures, bridging the gap between collated data and human understanding through natural language interaction.

**Fig. 4.**
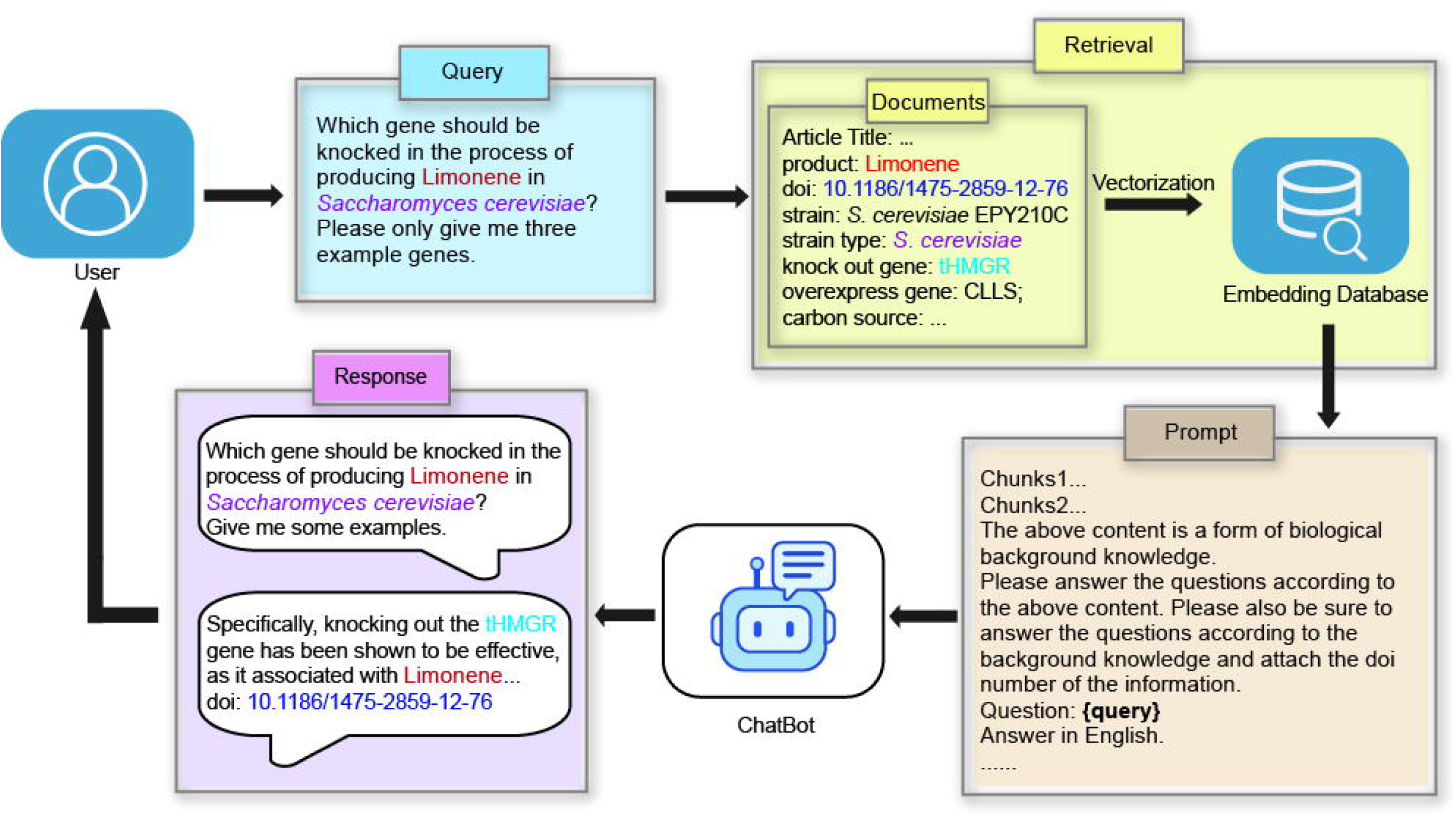
Workflow of the Metabolic Engineering Chatbot. This workflow integrates data from retrieved cell factory databases into an LLM-based dialogue system. The cell factory database is vectorized into an embedding database. The user’s initial query undergoes a similarity search within this embedding database and is then fed into the prompt instruction to generate a response. Follow-up queries are designed based on the conversation context to provide relevant replies. The doi for the reference literature was supplied to support the response.

### Metabolic engineering targets prediction model: D2Cell-pred

Metabolic engineering aims to optimize and redesign metabolic pathways in microorganisms or cells to boost the production of specific products or enable the synthesis of new ones^1^. However, predicting the impact of gene modifications, especially modification combinations, on product yield remains a challenge^7^. We developed D2Cell-pred, a hybrid model that combines mechanistic and deep learning approaches to predict outcomes for new cell factories, with the mechanistic component being a genome-scale metabolic model (GEM). D2Cell-pred takes as input the target product, the GEM structure, and a set of gene modifications, and outputs the predicted impact of these modifications on the product (Fig. 5a).

**Fig. 5.**
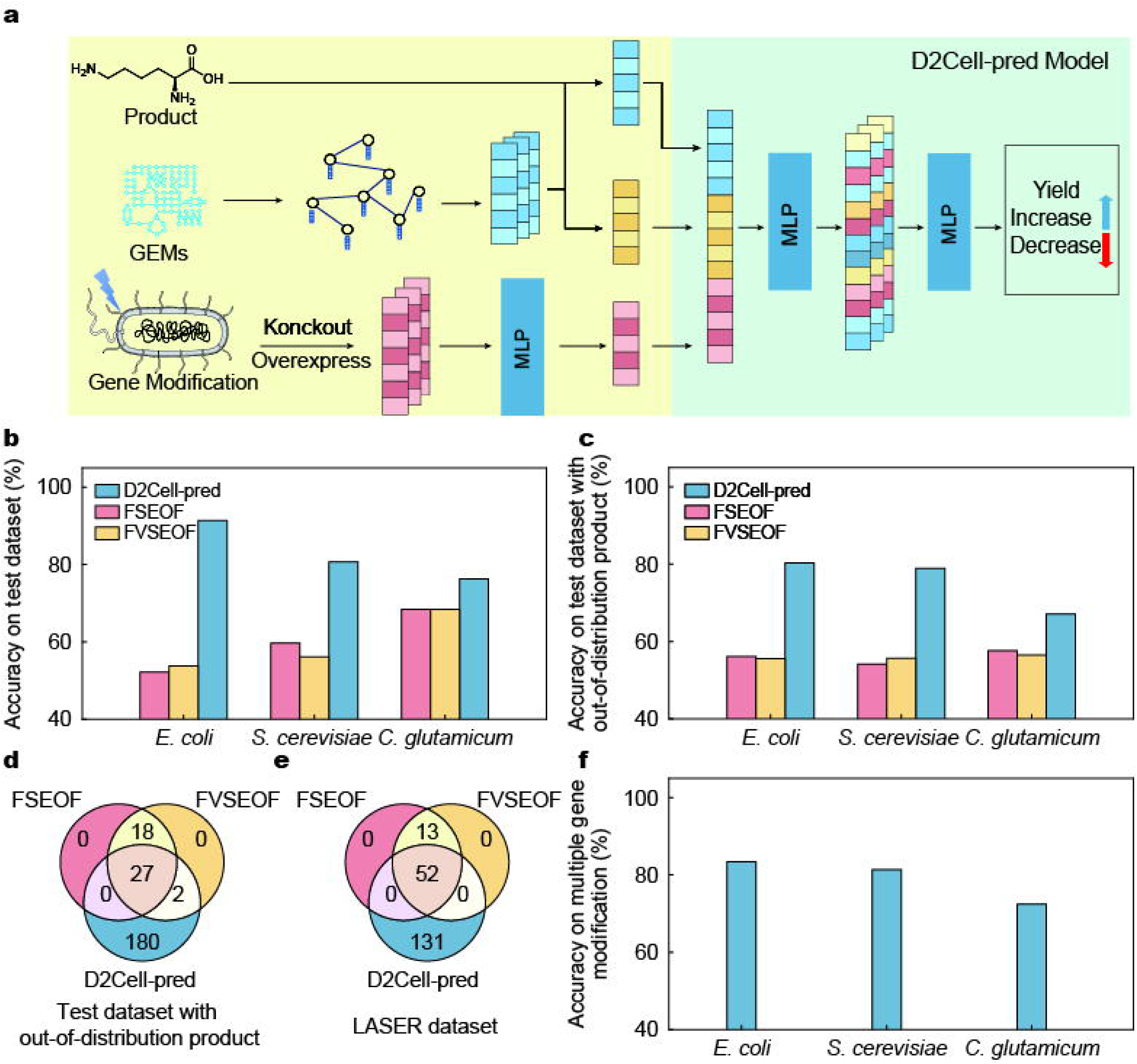
D2Cell-pred model architecture and performance. a) D2Cell-pred model, the approach developed for predicting impact of gene modifications on product yield. The input of the mode is the product, the GEM structure and the set of gene modifications. The output is the impact of the gene modifications towards the cell factory production. b) The accuracy of D2Cell-pred, FSEOF, and FVSEOF on test datasets for *E. coli*, *C. glutamicum* and *S. cerevisiae*. c) The accuracy of D2Cell-pred, FSEOF, and FVSEOF models on test datasets for out-of-distribution products for *E. coli*, *C. glutamicum* and *S. cerevisiae*. Out-of-distribution products means those products are not included in the training dataset. d, e) Numbers of correctly predicted target by D2Cell-pred, FSEOF, and FVSEOF methods on test datasets with out-of-distribution products and an independent LASER dataset for *E. coli*. f) The accuracy of multiple gene modification prediction on test datasets for *E. coli*, *C. glutamicum* and *S. cerevisiae*.

Since different organisms have different GEMs, we split the collected database by organisms, and for each organism, one model can be trained for cell factory prediction. We used three main model organisms *E. coli*, *S. cerevisiae* and *C. glutamicum* as examples to demonstrate the model capability. As for *E. coli*, the latest GEM iML1515^35^ was utilized and embedded in the cell factory model D2Cell-pred. For *S. cerevisiae* and *C. glutamicum*, we employed GEMs Yeast8^36^ and iCW773^37^, respectively. To supplement and expand our training dataset, we also employed the FSEOF method^2^ to generate simulated data. This approach can improve the model’s performance and generalization capability by creating additional data points^38^. The total dataset for *E. coli* consists of 8134 experimentally reported single- and double-gene modification data for 73 products and 11643 single gene modification data for 73 products simulated by GEM. The total dataset for *S. cerevisiae* consists of 2050 experimentally reported single- and double-gene modification data for 40 products and 18971 single gene modification data for 40 products simulated by GEM. And the total dataset for *C. glutamicum* consists of 1444 experimentally reported single- and double-gene modification data for 33 products and 3726 single- and double-gene modification data for 33 products simulated by GEM. The dataset was randomly partitioned into training (80%), validation (10%), and test (10%) sets. This random partitioning was repeated five times to confirm the deep learning model’s robustness (Supplementary Fig. 3a-c). Note that experimental entries with more than two gene modifications were excluded from this data set, serving as an independent test set to validate the model’s performance with multiple gene combinations in the further analysis.

### Prediction performance of D2Cell-pred for gene modification impact

The prediction performance of D2Cell-pred was systematically investigated (Supplementary File 4). Overall, D2Cell-pred achieved high accuracy with 0.914 for *E. coli*, 0.842 for *S. cerevisiae*, and 0.736 for *C. glutamicum* in the test dataset (Fig. 5b) with 10%-40% improvement compared to the commonly-used mechanistic algorithms such as FSEOF^2^ and its variant FVSEOF^39^. The slightly lower accuracy for *C. glutamicum* is likely due to the smaller amount of training data available for this organism. This overall strong performance demonstrates that D2Cell-pred effectively captures metabolic network topology and accurately predicts the impact of single- or double-gene modifications on target product production across different organisms. Even on the test dataset where products not included in the training dataset, D2Cell-pred still shows high accuracies in all three host organisms, with 0.804 for *E. coli*, 0.770 for *S. cerevisiae*, and 0.671 for *C. glutamicum* (Fig. 5c and Supplementary Fig. 4a-c), demonstrating that D2Cell-pred is capable of generalizing to out-of-distribution products. D2Cell-pred successfully identified 209 correct targets on the out-of-distribution test dataset, demonstrating superior performance compared to the FSEOF and the FVSEOF model, which identified only 45 and 47 targets, further showcasing D2Cell-pred’s ability to predict more targets overall than both FSEOF and FVSEOF in addition to the previously mentioned high accuracy (Fig. 5d). For example, D2Cell-pred identified several key targets for malate overproduction, including PTS system glucose-specific EIICB component (ptsG) target for reducing the glycolysis rate, the nicotinate phosphoribosyltransferase (pncB) target for expanding the NAD(H/+) pool, the fumarate reductase (frd) and fumarase (fumB) targets associated with the tricarboxylic acid cycle, and formate acetyltransferase (pflB). These targets for malate overproduction, which were reported in the literature^40^ but not included in the training dataset (and for which malate itself was not part of the training data), were not predicted by FSEOF and FVSEOF due to their ambiguous connection to malate production (Supplementary Fig. 5). These results suggest that D2Cell-pred can predict more comprehensive metabolic engineering strategies to optimize malate production in *E. coli*. The larger data set and higher proportion of experimentally validated information for *E. coli* provided significant qualitative and quantitative advantages, allowing the D2Cell-pred model to make more accurate predictions for *E. coli* compared to *S. cerevisiae* and *C. glutamicum*.

We assessed D2Cell-pred using the independent LASER dataset, which was not used during the model’s development, ensuring a fair benchmark. This dataset, containing only positive effective targets, includes 244 entries across 19 products from 39 studies on *E. coli*, all involving gene overexpression or knockout with glucose as the primary carbon source. We compared D2Cell-pred’s predictions with those from FSEOF and FVSEOF. D2Cell-pred outperformed the other methods, demonstrating superior accuracy for the majority of products (Supplementary Fig. 6), and accurately identifying 183 targets compared to 65 targets each identified by FSEOF and FVSEOF (Fig. 5e).

We also evaluated the ability of D2Cell-pred to predict the impact of multiple gene modifications (more than 2, can up to 12) (Supplementary File 5). To ensure the reliability of multi-gene modification predictions and to avoid the influence of single-gene modification training sets on the results, we constructed a multi-gene modification test set using products not seen in the dataset. Even though D2Cell-pred has not been trained on the more than two gene combination data, it demonstrated reasonable accuracy in this task, with over 80% accuracies on *E. coli* and *S. cerevisiae*, and 72% accuracy on *C. glutamicum* (Fig. 5f). We demonstrated that removing the GEM global feature module would decrease the D2Cell-pred performance, especially in the multi-gene combination tasks (Supplementary Fig. 7a-c). However, removing the global feature module from the model for *C. glutamicum* improved performance because the smaller training dataset made global features less relevant, introducing noise or irrelevant information. This is expected to be fixed with more data points collection.

## Discussion

Our research introduces a step-by-step approach to text mining by utilizing LLMs to extract extensive metabolic engineering knowledge from scientific literature. By harnessing the advanced capabilities of LLMs such as Qwen1.5 and GPT-4, we overcome the limitations of traditional manual data extraction and conventional natural language processing (NLP) models. The D2Cell-learn workflow refines text mining by breaking it into simpler, more manageable sub-tasks. This method significantly enhances the LLMs’ ability to accurately extract relevant data from research articles, improving both entity recognition and the precision of relational data extraction. Our comprehensive evaluation demonstrates how effectively these LLMs can be used to build a high-quality metabolic engineering knowledge database. The resulting cell factory database includes over 29006 entries, covering 751 strains and 1210 unique products. It provides extensive details on microbial organisms, products, and genetic modifications, making it an invaluable resource for researchers and training deep learning models^23^. To further enhance accessibility, we developed a chatbot using Retrieval-Augmented Generation (RAG) technology. This chatbot efficiently retrieves and presents information from the database, delivering precise and contextually relevant responses to user queries, and ensuring both accuracy and traceability. Overall, our study demonstrates the efficacy of LLMs as powerful tools in biological research, advancing the fields of text mining and knowledge extraction from scientific literature.

The availability of massive datasets enables us to tackle challenges previously hindered by data scarcity. For example, we developed a hybrid model for predicting engineering targets by combining GEM with graph neural networks. Trained on the extracted data, the model demonstrates superior performance and strong generalization compared to traditional methods^2, 39^, maintaining accuracy even for independent dataset^9^ and products that not included in training dataset. With more data available, we believe D2Cell-pred will be a powerful tool for cell factory design. Thus, future work can integrate more diverse literature sources such as patents and reviews. In addition, constrained by the current LLM performance and context length limit of LLMs, the methodological approach primarily concentrated on extracting information from textual content, failing to fully leverage the rich information embedded within figure illustrations and tabular data. However, these visual and tabular elements encapsulate invaluable knowledge pertaining to metabolic engineering. Effective utilization of this multimodal information is a direction for future exploration to improve the comprehensiveness and accuracy of knowledge extraction efforts^41^.

In conclusion, our work utilizes the powerful capability of LLMs to automatically extract metabolic engineering knowledge from existing literature to support applications in the field of metabolic engineering. Besides that, we also built a hybrid model for engineering target prediction and a conversational chatbot based on retrieval-enhanced generation technology to improve the accessibility to the dataset, demonstrating the potential of LLMs as assistant tools in the biological field.

## Methods

### D2Cell-learn and data extraction

We divided text mining D2Cell-learn into three parts: named entity recognition (NER), relation extraction (RE), and entity resolution and normalization (ER). Our data sources include abstracts of 10000 published research articles collected from Clarivate Web of Science, the National Library of Medicine PubMed and Scopus, along with 1340 open-access articles collected manually. For the open-access articles, we only use the abstract, results, and methods sections. Firstly, in the NER module, we split the articles into segments. We used the Qwen-Lora to extract ’strain ID’ and ’gene’ entities from segments of research papers along with corresponding prompts. The NER module returns the extracted information as a list of JSON objects. Although strain IDs often lack standardization across different studies—typically incorporating author initials and numbers such as “QW101” or “SA-B4”—we recognized their importance for tracking genetic modifications within the same study. Extracting strain ID entities in this step allows us to accurately trace genetic modification relationships and ensures the precision of subsequent relation extraction steps. In the RE module, we use the Qwen1.5-110B-Chat model to extract relationships between entities. Based on these relationships and the text in abstract, results and methods sections, we constructed prompts to extract experimental conditions for each entry, including temperature, vessel, culture time, medium composition, and carbon sources. Additionally, in the ER module, we obtained UniProt IDs and GeneBank IDs for the modified genes using UniProt’s Application Programming Interface (API)^42^. Moreover, a Python script was used to retrieve taxonomy IDs, standard names for each organism; and KEGG ENTRY for each product from the KEGG database^43^, from which we subsequently obtained the corresponding SMILES representations. For products without KEGG entries, we queried the PubChem compound database^44^, the largest database of chemical compound information, to get the corresponding information. The databases are hosted in an online database website (LINK). All prompts and codes used in this study can be found in the GitHub repository.

### Large language models

Due to the requirement of processing long tokens and high performance, we chose open-source large language model Qwen1.5^14^, capable of handling up to 32768 tokens. Considering performance and efficiency, we used the fine-tuned Qwen1.5-14B-Chat model for extracting strain ID and gene entities. For relation extraction tasks requiring long text inputs, due to hardware resource constraints precluding effective fine-tuning of models larger than 72B and the poor performance of smaller models in handling long texts, we used the un-fine-tuned Qwen1.5-110B-Chat model. The prompts for the text mining task are provided in the Supplementary Figure 8-14. We effectively fine-tuned the Qwen1.5-14B-Chat model using LoRA^45^, an approach that maintains most of the performance of full-parameter fine-tuning while significantly reducing computational requirements. We created a training dataset using Gemini Pro’s entity extraction results from 100 metabolic engineering research papers for fine-tuning Qwen1.5 model. To mitigate catastrophic forgetting when fine-tuning a large language model (LLM) and to enhance the extraction capabilities of the model, we augmented the training set with 20000 samples from the IEPile dataset^46^, which contains information on a variety of entities. The model was trained for 10 epochs with a learning rate of 5 × 10^-5^. Vllm^47^ was used to speed up inference for Qwen1.5-110B-Chat and Qwen1.5-14B-Chat. As baseline models, we selected widely used large language models, such as GPT-4 and Claude. However, due to the substantial cost of using GPT-4 and Claude to run the D2Cell text extraction method, we compared only the prompt without splitting the text extraction task against the D2Cell method. We used API keys to interact with ChatGPT and Claude, using model versions GPT-4o and Claude 2 Sonnet.

### Dataset preparation for D2Cell-pred

Since GEM has been widely used to predict metabolic engineering targets, including these simulations can expand the training dataset with a range of potential scenarios^2, 5^. By integrating both experimental and simulated datasets, the deep learning model aims to achieve comprehensive coverage of training data, leveraging GEMs’ predictive capabilities to enhance its ability to optimize and predict metabolic engineering outcomes effectively. We utilized Flux Scanning based on Enforced Objective Function (FSEOF)^2^ for simulated engineering target prediction.

Taken together, we generated the comprehensive datasets for *E. coli*, *S. cerevisiae*, and *C. glutamicum* including products, gene modifications such as knockouts and overexpression, and a binary classification indicating the impact of the gene modification on the product production (1 denotes enhancement, 0 denotes no or negative impact). The dataset for *E. coli*, *S. cerevisiae*, *and C. glutamicum* contain 19777, 21021, and 5170 entries, respectively. Literature predominantly documents the positive results, resulting in a dataset skewed towards positive outcomes. Therefore, we designated gene modifications opposing those enhancing product production in the literature as negative samples. Additionally, genes not predicted as targets by FSEOF were included in the negative dataset. In the datasets, positive samples account for 40% of the entries, while negative samples make up 60%. The datasets were split into training, testing, and validation sets with a ratio of 80%, 10%, and 10%, respectively. The comprehensive dataset was subsequently filtered to include only genes and products present in the GEMs. As for the GEM submodule, we utilized iML1515 model for *E. coli*^35^, Yeast8 for *S. cerevisiae*^36^ and iCW773 for *C. glutamicum*^37^.

### Comparison with commonly used target prediction algorithms FSEOF and FVSEOF

We employed the FSEOF^2^ and FVSEOF^39^ algorithms together with genome-scale metabolic models Yeast8^36^, iML1515^35^, and iCW773^37^ to identify overexpression and knockdown targets for various products in *S. cerevisiae*, *E. coli*, and *C. glutamicum*, respectively.

For the FSEOF algorithm, we utilized pFBA^48^ to simulate growth on glucose across a range of biomass yields, from zero to maximum, distributed at uniform intervals. All unused carbon for biomass accumulations was directed towards different target product production. Reactions exhibiting a positive correlation between flux and product synthesis were designated as overexpression targets. Conversely, reactions showing a negative correlation with product synthesis and not essential for growth were identified as knockout targets.

For FVSEOF, the upper and lower limits of each metabolic reaction were determined by flux variability analysis at each biomass yield (ranging from 0 to maximum biomass yield, distributed at uniform intervals). Reactions for which both the upper and lower flux bounds positively correlated with product synthesis were classified as overexpression targets. Similarly, reactions where both flux bounds negatively correlated with product synthesis, while not being critical for growth, were designated as knockout targets.

### Construction of the D2Cell-pred

D2Cell-pred integrates a GEM as a mechanistic submodule with a deep learning architecture. Since a GEM can be effectively represented as a graph, with nodes denoting metabolites and edges representing metabolic reactions^49^, the model utilizes a graph neural network (GNN) structure to represent GEM. The GNN captures features of metabolite embeddings within the stoichiometric matrix. Genetic modifications are represented by embeddings that are refined during the training process to capture their key characteristics. The embedding information for each modification in a gene modification set is combined with the target product embeddings. The product embedding is shared with the same metabolite in the GEM structure, enable the deep learning model to understand the product’s relationship within the network.

D2Cell-pred was designed to predict the impact of genetic modifications on product production by combining features from the products, genetic modifications, and the metabolic network. Hyperparameters used for model training are provided in Supplementary Table 2. D2Cell-pred utilized learnable embeddings to represent each product 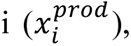 rather than using scalar representation, allowing the model to capture relative heterogeneity among products. To capture relationships between different products, we applied a GNN on the metabolic network graph (*G_GEMs_*), where nodes represent products and edges represent reactions. Given this metabolic network graph, the GNN can iteratively update the vector representation of each metabolite by propagating and transforming information from its neighboring metabolites through non-linear functions within the neural network layers. The final output of the GNN module is a set of new feature vectors 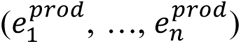 that represent the metabolites, encoding information from both the products and the metabolic network topology. These features fully consider the associations between products and can accurately represent their roles and functions within the metabolic network, shown in equation (1).

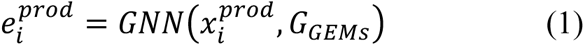

Besides the node feature, we applied another GNN was applied to the metabolic network to extract graph-level features *g^GEMs^*. Unlike node features that primarily capture local information around connected nodes, global features can reflect the overall structure and patterns of the graph. Similar to product feature extraction, we used learnable embedding representation for all gene modifications. Given a set of gene modifications (*M*_1_, …, *M_k_*), the model extracts the embeddings of the input gene 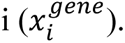 We employ a combination strategy to integrate the embeddings of multiple gene modifications, where the embeddings are aggregated through a summation operation, and the resulting representation is subsequently processed by a MLP, shown in equation (2),

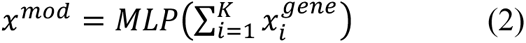

We used MLP modules to capture the relationship between different features. We concatenated the products feature, gene modifications feature, and the metabolic network graph feature, and feed them into a MLP module to obtain a feature vector that encapsulates the impact of gene modifications. This feature vector was subsequently fed into a multi-layer perceptron (MLP) to map the features to the final classification output, shown in equation (3) and (4).

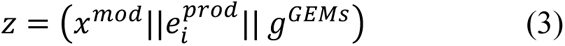

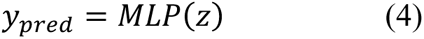

Where || refers to the operation of vector concatenation; *y_pred_* is the predict label.

During the training process, we first shuffled all the data, and then randomly split it into a training set, validation set, and test set with a ratio of 80%:10%:10%. Given a set of products, a metabolic network graph, a set of gene modifications, and the corresponding class labels, the objective of the training process was to minimize the cross-entropy loss function. The best model was selected based on the minimum cross-entropy, as shown in equation (5).

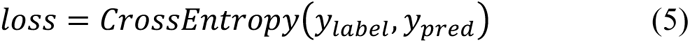

Where *y_label_* is the ground truth label.

### ChatBot-Retrieval-Augmented LLMs

We implemented a RAG approach to enhance the performance of LLMs as chatbot. To achieve this objective, we utilized the BGE-M3 embedding model developed by BAAI, a robust multilingual text embedding model^50^, to transform the original tabular data into high-dimensional vector representations. Our RAG system architecture was implemented using the LangChain framework, an open-source platform designed for developing LLM applications, which offers a comprehensive suite of components and tools facilitating the construction of sophisticated AI applications. In the retrieval phase, we employed a semantic similarity search methodology. Specifically, for each user input, we selected the top 5 most semantically similar tabular data entries as relevant context. Upon completion of the retrieval phase, we integrated the retrieved tabular data with the user input to construct a comprehensive prompt. This comprehensive prompt was subsequently input into the Qwen1.5-110B-Chat model, for processing and generation of the final response.

## Data availability

The yeast consensus genome-scale model version 8.0.1^36^, *E. coli* genome-scale model iML1515^35^, *C. glutamicum* genome-scale model iCW773^37^ were used for simulations. Clarivate Web of Science (https://mjl.clarivate.com/home), National Library of Medicine PubMed (https://pubmed.ncbi.nlm.nih.gov/) and Scopus (https://www.scopus.com/sources) database were used to collect articles abstract. LASER database (https://bitbucket.org/jdwinkler/laser_release/src/master/) was used to evaluate the model performance. IEPile dataset (https://github.com/zjunlp/IEPile) was used to fine-tune Qwen model. UniProt database (https://www.uniprot.org/) and NCBI database (https://www.ncbi.nlm.nih.gov/) were used to obtain UniProt IDs and GeneBank IDs for the modified genes. KEGG database (https://www.genome.jp/kegg/) and PubChem database (https://pubchem.ncbi.nlm.nih.gov/) were used to obtain SMILES and KEGG ENTRY of product. All data used in the paper can be found in the GitHub repository: https://github.com/LiLabTsinghua/D2Cell.

## Code availability

In order to facilitate additional utilization, we have made available all of the codes and thorough instructions in our GitHub repository located at https://github.com/LiLabTsinghua/D2Cell.

## Author Contributions

F.L., J.N., Y.C., X.L. and Z.L. designed the research. X.L., Z.G. and Z.L. performed the research. X. L., Z. L., Z. G., 2, Z. L., K. W., J. L., Y. Z., L. L., M. S., H. T., Y. C., T. Y., J. N. and F. L. analyzed the data. F.L. and X.L. wrote the paper. K.W. performed engineering target prediction. Y.H. developed the website. All authors approved the final paper.

