## Supplementary Fig. for "Leveraging large language models for metabolic engineering design"

Supplementary Table 1. NER Performance of different LLMs

| NER task | Model | Precision | Recall | F1 score |
|---|---|---|---|---|
| | Qwen1.5-110B-Chat | 0.644 | 0.889 | 0.747 |
| | Qwen1.5-14B-Chat | 0.503 | 0.739 | 0.599 |
| | Llama3-8B-Instruct | 0.576 | **0.892** | 0.700 |
| Gene | Gemini Pro | 0.817 | 0.681 | 0.743 |
| | Claude3 | 0.611 | 0.878 | 0.720 |
| | GPT4 | 0.695 | 0.890 | 0.780 |
| | Llama-Lora | 0.726 | 0.875 | 0.794 |
| | Qwen-Lora | **0.825** | 0.868 | **0.846** |
| | Qwen1.5-110B-Chat | 0.620 | **0.927** | 0.743 |
| | Qwen1.5-14B-Chat | 0.623 | 0.748 | 0.680 |
| | Llama3-8B-Instruct | 0.566 | 0.894 | 0.693 |
| Strain ID | Gemini Pro | 0.856 | 0.760 | 0.805 |
| | Claude3 | 0.761 | 0.681 | 0.718 |
| | GPT4 | 0.743 | 0.835 | 0.713 |
| | Llama-Lora | 0.781 | 0.769 | 0.774 |
| | Qwen-Lora | **0.894** | 0.857 | **0.876** |

Supplementary Table 2. Hyperparameters used in D2Cell-Pred

| Model organism | Epochs | Learning rate | Batch size | Optimizer | Number of metabolites embedding |
|---|---|---|---|---|---|
| *E. coli* | 500 | 5e-5 | 16 | Adam | 2000 |
| *S. cerevisiae* | 500 | 5e-5 | 16 | Adam | 3000 |
| *C. glutamicum* | 500 | 5e-5 | 16 | Adam | 2000 |

*E. coli*: *Escherichia coli*, *S. cerevisiae*: *Saccharomyces cerevisiae*, *C. glutamicum*: *Corynebacterium glutamicum*

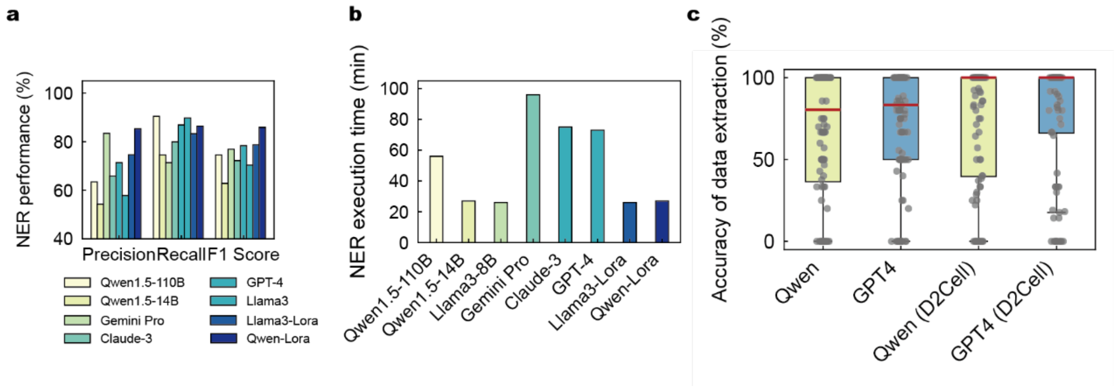


**Supplementary Fig. 1. Data extraction performance on test data.** a) Overall precision, recall and F1 scores of different LLMs on the NER task. b) Execution time of different LLMs on the NER task for 873 text segments. c) Paper-based accuracy of the data extracted from 100 cell factory articles in the test set. Each dot represents one paper. In box plot, the central band represents the median value, the box represents the upper and lower quartiles, and the whiskers extend up to 1.5 times the interquartile range beyond the box range.

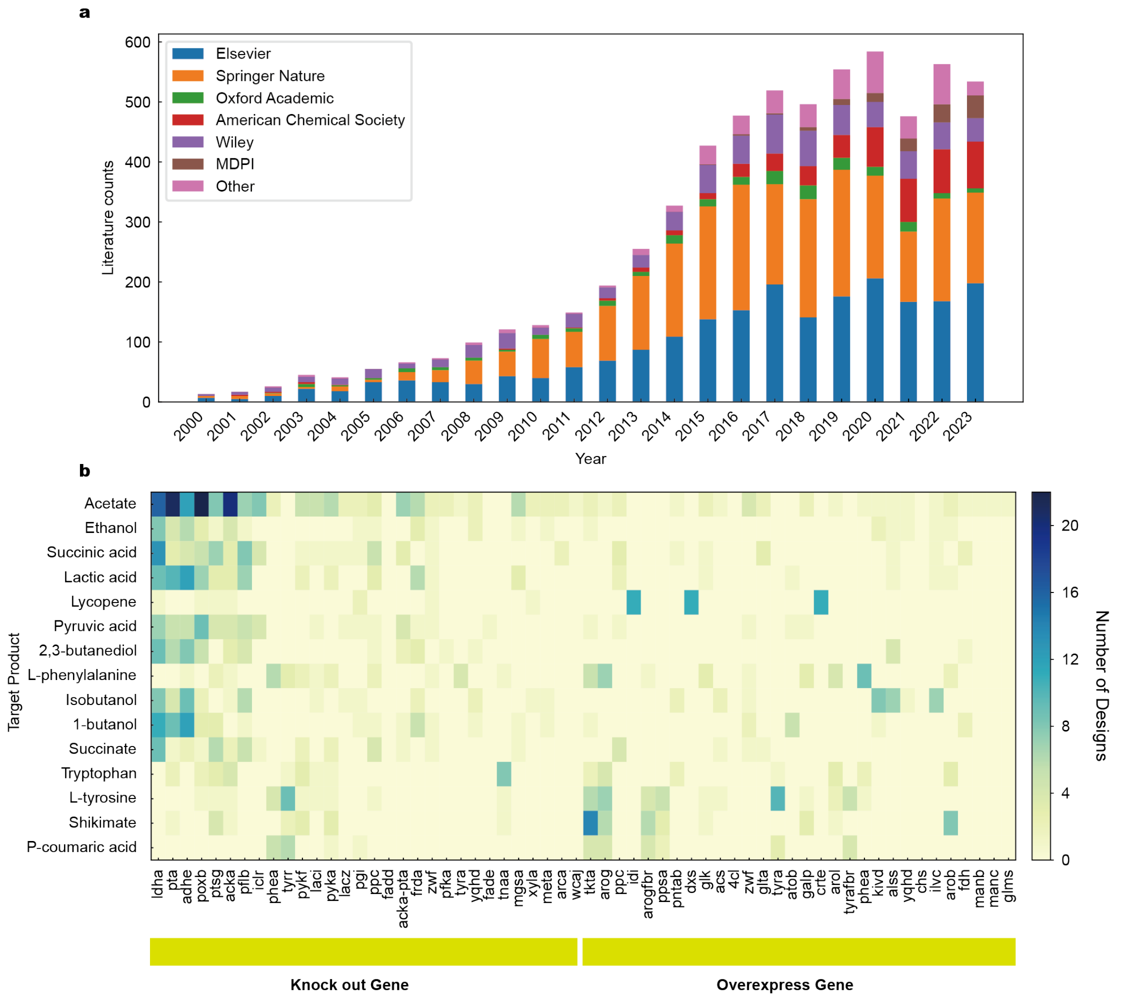


**Supplementary Fig. 2. Statistical analysis of cell factory database.** a) Number and distribution of articles in the database by publication year (2000-2023) for the top six publishers' group in the database. b) The number of studies reporting different gene modifications for each product. Top 30 common genes were displayed for 15 products in *E. coli*.

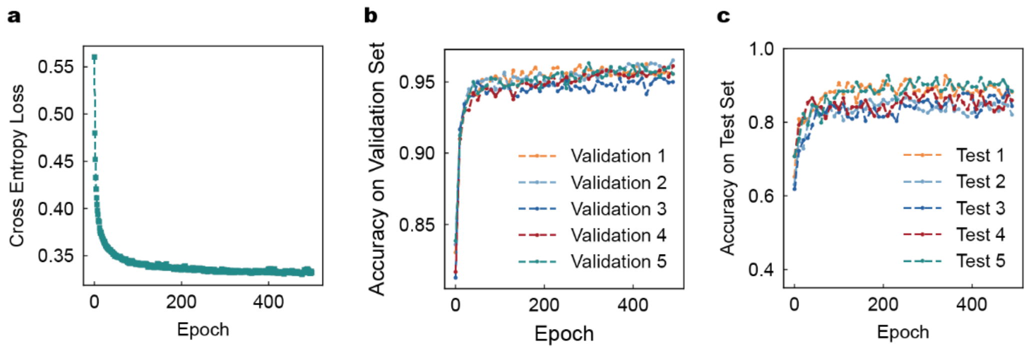


**Supplementary Fig. 3. Training performance of the D2Cell-pred on *E. coli*.** a) Loss in training D2Cell-pred. Performance of the D2Cell-pred on b) validation dataset and c) test dataset by running five times of random splitting.

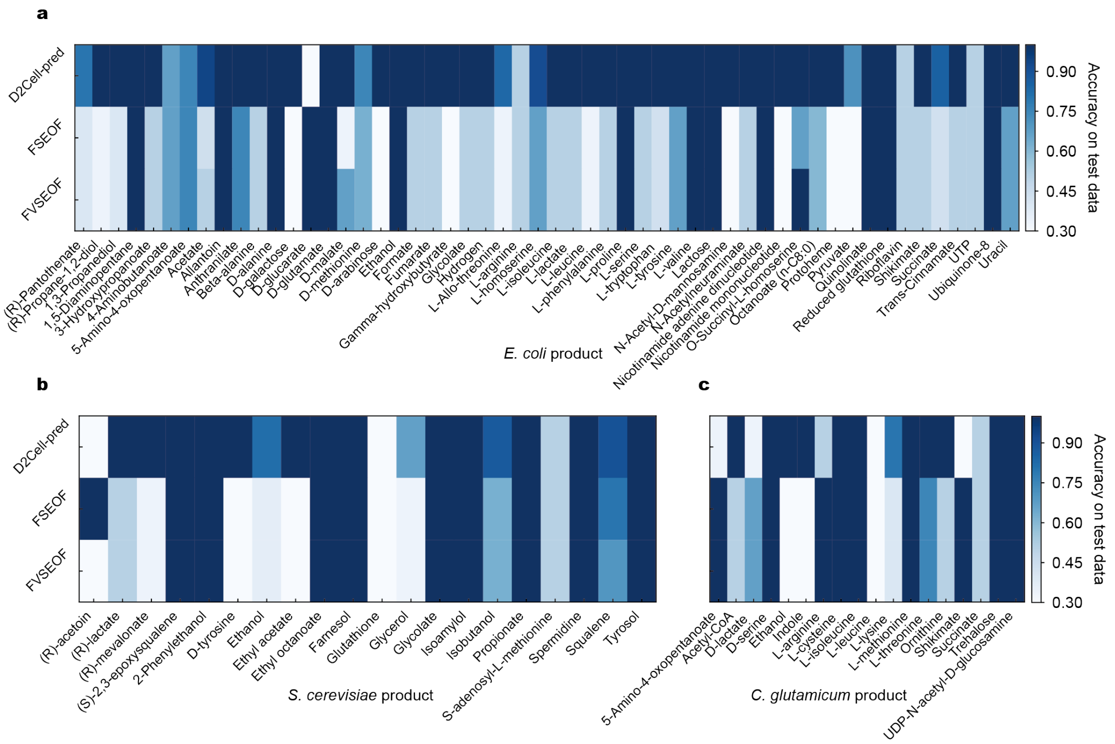


**Supplementary Fig. 4. Comparison of prediction accuracy for various products using the D2Cell-pred and baseline models on test datasets** for a) *E. coli*, b) *S. cerevisiae*, and c) *C. glutamicum*. *E. coli*: *Escherichia coli*, *S. cerevisiae*: *Saccharomyces cerevisiae*, *C. glutamicum*: *Corynebacterium glutamicum*.

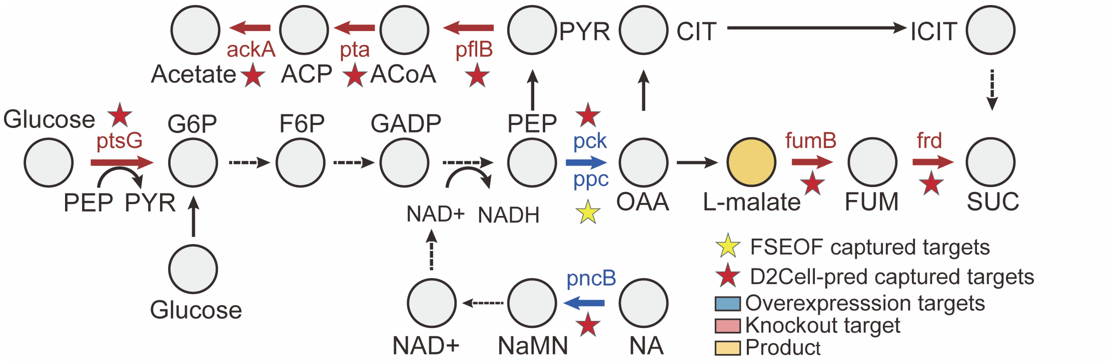


**Supplementary Fig. 5. Predicted targets for L-malate production in *E. coli*.** This example study from the out-of-distribution products dataset demonstrates the use of the D2Cell-pred model for L-malate production, highlighting overexpression and knockout targets. Blue arrows and text denote genes and reactions associated with overexpression, while red arrows and text indicate knockout genes and reactions. Red stars mark overexpression or knockout genes and reactions not predicted by D2Cell-pred, yellow stars represent those predicted by FSEOF. Dotted arrows show multiple reactions. FUM: Fumarate; SUC: Succinate; CIT: Citrate; ICIT: Isocitrate; PEP: Phosphoenolpyruvate; OAA: Oxaloacetate; ACoA: Acetyl-coenzyme A; ACP: Acetyl phosphate; NA: Nicotinic acid; NaMN: Nicotinic acid mononucleotide; G6P: Glucose-6-P; F6P: Fructose-6-P; GADP: Glyceraldehyed 3-P; PYR: Pyruvate; ppc: Phosphoenolpyruvate carboxylase; pck: Phosphoenolpyruvate carboxykinase; fumB: Fumarase; ackA: Acetate kinase; ptsG: Phosphotransferase system; pflB: Pyruvate-formate lyase; pncB: NA phosphoribosyltransferase; pta: Phosphate acetyltransferase; frd: Fumarate reductase.

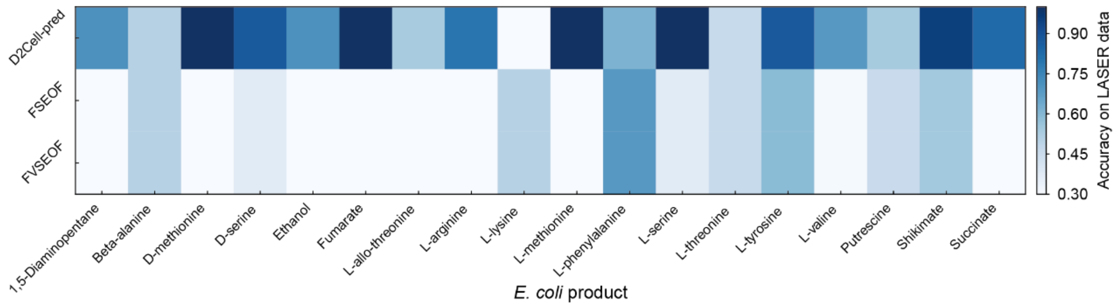


**Supplementary Fig. 6. Comparison of prediction accuracy for various products using the D2Cell-pred and baseline models on LASER datasets.**

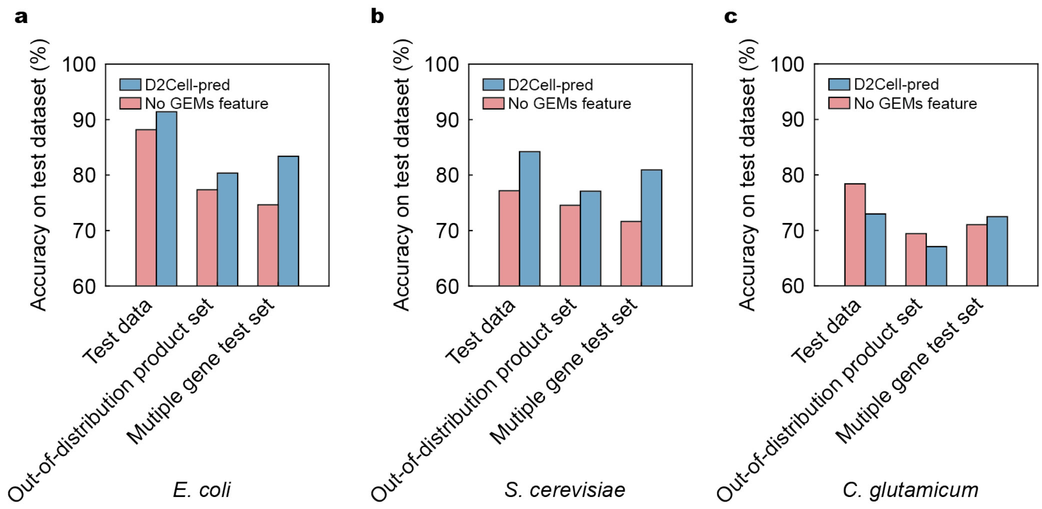


**Supplementary Fig. 7. Model ablation study on test dataset.** Ablation studies were performed on D2Cell-pred models trained using a) *E. coli*, b) *S. cerevisiae* and c) *C. glutamicum*. The test dataset with out-of-distribution product refers to the test dataset filtered to only include products that were not included in the training set. The 'No GEMs Feature' condition removes the GEMs global feature. This study evaluates the importance of different components in D2Cell-pred and their impact on accuracy using the test dataset.

**Gene extraction prompt:**
You need to read the following input text carefully.
Your task is to extract all gene names mentioned in the input text.

**{Input text}**

You should provide the answer in JSON array format as the exmaple
{"gene name": []}.

---

**Input text:**
In E. coli, genes xylA and xylB encoded xylose isomerase and xylulose kinase, respectively. Deletion of xylA and xylB and overexpression the first two genes of Weimberg pathway in E. coli increased xylonate production from xylose (Cao et al. 2013; Liu et al. 2012a). In this study, genes xylA and xylB were deleted in WWZ04 and the plasmid pairs pWZt7-g2/pWZt7-xyl and pWZtac-g2/pWZtac-xyl were transformed into the sextuple deletion mutant WWZ06 (Fig. 4).

---

**Output:**
{"gene name": ["xylA", "xylB"]}.

**Supplementary Fig. 8 An illustration of prompt engineering for gene extraction in NER task.**

**Strain extraction prompt:**
You need to read the following input text carefully.
Your task is to extract all strain names mentioned in the following input text.

**{Input text}**

You should provide the answer in JSON array format as the exmaple {"strain name": []}.

---

**Input text:**
This was because OptForce suggested to upregulate particular reactions in the mevalonate pathway (Additional file 1: Table S5) for GGPP overproduction, and these were already improved in our engineered strain, LRS6. As our engineered yeast strains contained additional genes driven by galactose-inducible promoters, the parent strain KM1, was not able to induce the integrated mevalonate pathway genes using glucose as sole carbon source. Therefore, we first deleted the GAL80 gene yielding with EJ1 strain (Additional file 1: Table S4) to allow glucose utilization as Gal80 protein inhibits the transcription of the galactose inducible genes in the absence of galactose [48].

---

**Output:**
{"strain name": ["LRS6", "KM1", "EJ1"]}.

**Supplementary Fig. 9 An illustration of prompt engineering for strain ID extraction in NER task.**

**Product extraction prompt:**
You need to read the following text carefully and answer my questions.

**{Input text}**

Your task is to extract all products produced by the strain **{Input strain ID}** from the above input text. You should provide the answer in JSON format as the exmaple. Here is an example of what the answer should look like: {"product": []}.
If there is no product in the above text, you should answer {}.

---

**Input text:**
Among the typical 2,3-BD producers, B. subtilis 168 produces (2R,3R)-2,3-BD as the major product (Ji et al., 2011, Zhang et al., 2013). B. licheniformis 10-1-A produces (2R,3R)-2,3-BD and meso-2,3-BD with a ratio of nearly 1:1 (Li et al., 2013). K. pneumoniae and Enterobacter cloacae produce meso-2,3-BD and (2S,3S)-2,3-BD as the major products (Ji et al., 2011, Wang et al., 2012).

---

**Input strain ID:**
B. licheniformis 10-1-A

---

**Output:**
{"strain": ["B. licheniformis 10-1-A"], "product": ["(2R,3R)-2,3-BD", "meso-2,3-BD"]}.

**Supplementary Fig. 10 An illustration of prompt engineering for product extraction in RE Task.**

**Product titer extraction prompt:**
You need to read the following text carefully and answer my questions.

**{Input text}**

The above input text contains information about the different products and product yields and product titers.Your task is to extract all product titers and product yields for the product **{Input product}** from the input text above, without missing any product yield or titer data. Product yield must be in g/g or can be converted to g/g. Product titer must be in g/l or can be converted to g/l. Output must contain the unit. You should provide the answer in JSON format as the exmaple {"product": "","product titers":[""], "product yields":[""]}. If no product titers or yields appear in the article, answer {}.

---

**Input text:**
The combined rational engineering increases 3-HP production from 0.14 g/L to 11.25 g/L in shake flask using 20 g/L glucose, approaching the maximum theoretical yield with concurrent biomass formation. The engineered yeast forms the basis for commercialization of bio-acrylic acid, while our CO2 fixation strategies pave the way for CO2 being used as the sole carbon source.

---

**Input product:**
3-HP

---

**Output:**
{"product": "3-HP", "product titers":["0.14 g/L", "11.25 g/L"], "product yields":[""]}.

**Supplementary Fig. 11 An illustration of prompt engineering for product titer extraction in RE Task.**

**Strain and gene relationship extraction prompt:**
You need to read the following text carefully and answer my questions.

**{Input text}**

According to the input text above, you should find the genetic modifications corresponding to the strain, including which genes the strain knocks out, which genes are overexpressed, and which genes are heterologous gene.
**{Input strain ID}**
**{Input gene}**
The above are the genes and strain names you need to consider. You will need to fill out the form below based on the information you have been given. You need to output the answer in markdown format.
| strain name | knockout gene | overexpress gene | heterologous gene | parent strain |

---

**Input text:**
As our engineered yeast strains contained additional genes driven by galactose-inducible promoters, the parent strain KM1, was not able to induce the integrated mevalonate pathway genes using glucose as sole carbon source. Therefore, we first deleted the GAL80 gene yielding with EJ1 strain (Additional file 1: Table S4) to allow glucose utilization as Gal80 protein inhibits the transcription of the galactose inducible genes in the absence of galactose [48].

---

**Input strain ID:**
KM1, EJ1

---

**Input gene:**
GAL80

---

**Output:**

| strain name | knockout gene | overexpress gene | heterologous gene | parent strain |
|---|---|---|---|---|
| EJ1 | GAL80 | | | |

**Supplementary Fig. 12 An illustration of prompt engineering for strain and gene relationship extraction in RE Task.**

**Strain and product relationship extraction prompt:**
You need to read the following text carefully and answer my questions.

**{Input text}**

Between the three single quotes is the input text. The follwing list is the name of the strains all you need to consider.
The strain names mentioned in the text include: **{Input strain ID}**.
Based on the information detailed in the input text, which strain produces **{Input product}**?
You should provide the answer in JSON format as the exmaple {"strain": ""}.

---

**Input text:**
Here, we used systemic metabolic engineering strategies to debottleneck the 2,3-BDO production in Enterobacter aerogenes. Firstly, the pyruvate metabolic network was reconstructed by deleting genes for by-product synthesis to improve the flux toward 2,3-BDO synthesis, which resulted in a 90% increase of the product titer. Secondly, the 2,3-BDO productivity of the IAM1183-LPCT/D was increased by 55% due to the heterologous expression of DR1558 which boosted cell resistance to abiotic stress. Thirdly, carbon sources were optimized to further improve the yield of target products. The IAM1183-LPCT/D showed the highest titer of 2,3-BDO from sucrose, 20% higher than that from glucose, and the yield of 2,3-BDO reached 0.49 g/g. Finally, the titer of 2,3-BDO of IAM1183-LPCT/D in a 5-L fermenter reached 22.93 g/L, 85% higher than the wild-type strain, and the titer of by-products except ethanol was very low.

---

**Input strain ID:**
IAM1183-LPCT/D

---

**Input product:**
0.49 g/g 2,3-BDO

---

**Output:**
{"strain": "IAM1183-LPCT/D"}

**Supplementary Fig. 13 An illustration of prompt engineering for strain and product relationship extraction in RE Task.**

**Cultivation conditions extraction prompt:**
You need to read the following text carefully and fill up the given form according to the article.

**{Input text}**

Between the three single quotes is the input text. The table below contains the strains and products of the strains as well as the product titers. You need to fill in what experimental conditions each product titer was produced under. You only need to fill in the blanks in the form I have given you. The output is a table in markdown format.

**{Input table}**

---

**Input text:**
All microorganisms were grown at 30 \u00b0C in LBB broth (Luria\u2013Bertani broth with 18.5 g/L brain heart infusion) or on LBB agar plates. LB consists of 10 g/L tryptone, 5 g/L yeast extract, and 10 g/L NaCl). The seed medium contained the following components in units of g/L: glucose, 25.0; corn syrup, 20.0; KH2PO4, 1.0; (NH4) 2SO4, 0.5; and urea, 1.25; and was adjusted to a final pH of 7.0.

---

**Input table:**

| strain | product | product titer | medium | Carbon source | Carbon source concentration | Vessel and feed mode | pH | Time | temperature |
|---|---|---|---|---|---|---|---|---|---|
| ATCC 13032 | GlcNAc | 0.5 g/L | | | | | | | |

---

**Output:**

| strain | product | product titer | medium | Carbon source | Carbon source concentration | Vessel and feed mode | pH | Time | temperature |
|---|---|---|---|---|---|---|---|---|---|
| ATCC 13032 | GlcNAc | 0.5 g/L | LBB | Glucose | Not specified | Batch | 7.0 | 72 h | 30 \u00b0C |

**Supplementary Fig. 14 An illustration of prompt engineering for cultivation condition extraction in RE Task.**
